# CpG-creating Mutations are Costly in Many Human Viruses

**DOI:** 10.1101/702175

**Authors:** Victoria R. Caudill, Sarina Qin, Ryan Winstead, Jasmeen Kaur, Kaho Tisthammer, E. Geo Pineda, Oana Carja, Rosalind M Eggo, Katia Koelle, Katrina Lythgoe, Scott Roy, Nicole Allen, Milo Aviles, Brittany A. Baker, William Bauer, Shannel Bermudez, Corey Carlson, Francisca L. Catalan, Angeline Katia Chemel, Dwayne Evans, Natalie Fiutek, Emily Fryer, Samuel Melvin Goodfellow, Mordecai Hecht, Kellen Hopp, E. Deshawn Hopson, Amirhossein Jaberi, Christen Kinney, Derek Lao, Adrienne Le, Jacky Lo, Alejandro G. Lopez, Andrea López, Fernando G Lorenzo, Gordon T Luu, Andrew R Mahoney, Rebecca L. Melton, Gabriela Do Nascimento, Anjani Pradhananga, Nicole S. Rodrigues, Annie Shieh, Jasmine Sims, Rima Singh, Hasan Sulaeman, Ricky Thu, Krystal Tran, Livia Tran, Elizabeth J. Winters, Albert Wong, Pleuni S. Pennings

## Abstract

Mutations can occur throughout the virus genome and may be beneficial or deleterious. We are interested in mutations that yield a C next to a G, producing CpG sites. CpG sites are rare in eukaryotic and viral genomes. For the eukaryotes, it is thought that CpG sites are rare because they are prone to mutation when methylated. In viruses, we know less about why CpG sites are rare. A previous study in HIV suggested that CpG-creating transition mutations are more costly that similar non-CpG-creating mutations. To determine if this is the case in other viruses, we analyzed the allele frequencies of CpG-creating and non-CpG-creating mutations across various strains, subtypes, and genes of viruses using existing data obtained from Genbank, HIV Databases, and Virus Pathogen Resource. Our results suggest that CpG sites are costly for most viruses. By understanding the cost of CpG sites, we can obtain further insights into the evolution and adaptation of viruses.

## Introduction

Viruses cause a multitude of diseases such as AIDS, Dengue Fever, Polio, Hepatitis, and the flu. Due to their fast replication, large population sizes and high mutation rates, viruses are able to quickly adapt to new environments (Cuevas et al., 2015). The ability of viruses to adapt quickly is seen in drug resistance evolution in HIV and HCV, immune escape in influenza and vaccine-derived polio outbreaks. High mutation rates may also lead to a high mutational load, since a large proportion of mutations are costly to the virus. In fact, experimental work has shown that most mutations are deleterious for viruses, with a select few being neutral or beneficial (Sanjuán et al., 2004; Duffy, 2018). Costs of mutations can also be studied using phylogenetic approaches (Stern et al., 2007) and within-host diversity data (Zanini et al., 2017; Theys et al., 2018).

Several different types of studies found evidence that CpG sites are costly for viruses. A CpG site refers to an occurrence of a nucleotide C followed by G in the 5’ to 3’ direction. Studies of viral genomic sequences found that CpG sites were underrepresented in almost all small viruses tested (Karlin and Cardon, 1994). In 2009, Burns et al. found that CpG sites significantly decreased replicative fitness of polio viruses in vitro, while an increased CG content in itself had little to no effect on the virus’s overall fitness. Stern et al. (2017) showed that CpG sites in the polio vaccine were often mutated in vaccine-derived polio outbreaks, indicating a direct cost of CpG sites in polio in vivo. In 2018, a study by Theys et al. found that in HIV, transition mutations resulting in CpG sites, were twice as costly as -otherwise similar- non-CpG-creating mutations, thereby revealing that CpG mutations have a cost within the host.

It is not entirely clear why CpG sites are costly, but it is likely, at least in part, because the mammalian immune system uses CpG sites to recognize foreign genetic material (Murphy and Weaver, 2016). Recently it was shown that ZAP proteins, which inhibit the proliferation of most RNA viruses, were more effective when the CpG sites were common (Takata et al., 2017).

A previous paper from our group (Theys et al., 2018) focused on the cost of CpG-creating mutations in HIV. Here we expanded our scope to encompass an array of human viruses, including Dengue, Influenza, Entero, Herpes, Hepatitis B C, and Polio. We focused on human viruses with a sufficient number of available sequences in Genbank, HIV Databases, or The Virus Pathogen Resource (VPR). Unlike in the Theys et al. (2018) paper, we focus on population-wide data (one sequence per patient) as opposed to within-patient data. The main assumption for this study is that when CpG-creating mutations come with a cost (either within hosts or at the transmission stage), we expect them to occur at lower frequencies in the population-wide sample compared to non-CpG-creating mutations. Since the types of mutations we consider (CpG-creating and non-CpG-creating) all occur on the same species-wide genealogy, we consider any significant differences in frequencies to be likely the result of a difference in cost. For a second analysis, we assume that the average frequency of mutations is inversely proportional to the cost of the mutations. This is likely an oversimplification, but it allows us to quantify the effect size we observe.

Depending on data availability, either individual genes or whole genomes were used. We found that CpG sites are costly in most viruses, though the effect is much stronger in some viruses (e.g., HIV) than others (e.g., HCV).

## Results

We collected 43 viral datasets from online sources (Genbank, HIV Dataset, VPR), each of which is a group of viral sequences of the same species, subtype and gene (see Table 2). Each sequence in a dataset came from an individual host from various parts of the world. The mean number of sequences in a dataset is 2,655, median 902, with a maximum at 24,005 and a minimum at 41. The mean number of nucleotides for each sequence is 3,344, median 1,617, with a maximum of 15,471 and a minimum of 181. We established a minimum cut off of 60,000 data points per dataset (number of nucleotides ∗ number of sequences), viruses or genes with less data available were not included.

We use the following approach. We assume that mutations occur at random, but are then subject to selection and drift. Selection and drift can act within hosts or at the transmission stage. For most mutations, selection will act to purge the mutations from the viral population (within-host population or the global population). Whether within-host or between-host effects are more important is not clear for most viruses, but either way, we expect that more deleterious mutations are less likely to be observed often, and more benign mutations will be observed more often. The main focus of our paper is to determine whether CpG-creating mutations are observed less often in each of the 43 datasets than (otherwise similar) non-CpG-creating mutations. We focus on A→G and T→C mutations, because transition mutations are more common in viruses than transversion mutations and only these transition mutations can create CpG sites.

To check whether our approach was sound, in principle, and whether there was sufficient power to asses the cost of CpG-creating mutations, we first tested whether synonymous mutations were observed at higher frequencies than non-synonymous mutations using the non-parametric Wilcoxon test. All tests are one-tailed, because we expect synonymous mutations to occur at a higher frequency than non-synonymous mutations. To make our approach for non-synonymous sites as similar as possible to our approach for CpG-creating mutations, we also focus solely on A→G and T→C mutations. We observed a significant difference between the frequencies of synonymous mutations and non-synonymous mutations for 35 of the 43 datasets analyzed (81.4 %) (table 1).

**Table 1:** The number of data sets for which the Wilcoxon test was significant (percentages in parentheses) indicating that non-CpG-creating mutations were observed at higher frequencies than, otherwise similar, CpG-creating mutations.

| Comparisons | A → G | T → C |
| --- | --- | --- |
| Synonymous: CpG vs non-CpG | 32 (74.5%) | 31 (72.1%) |
| Non-synonymous: CpG vs non-CpG | 11 (25.6%) | 13 (30.2%) |
| Synonymous vs Non-synonymous | 35 (81.4%) | 35 (81.4%) |

**Table 2:**
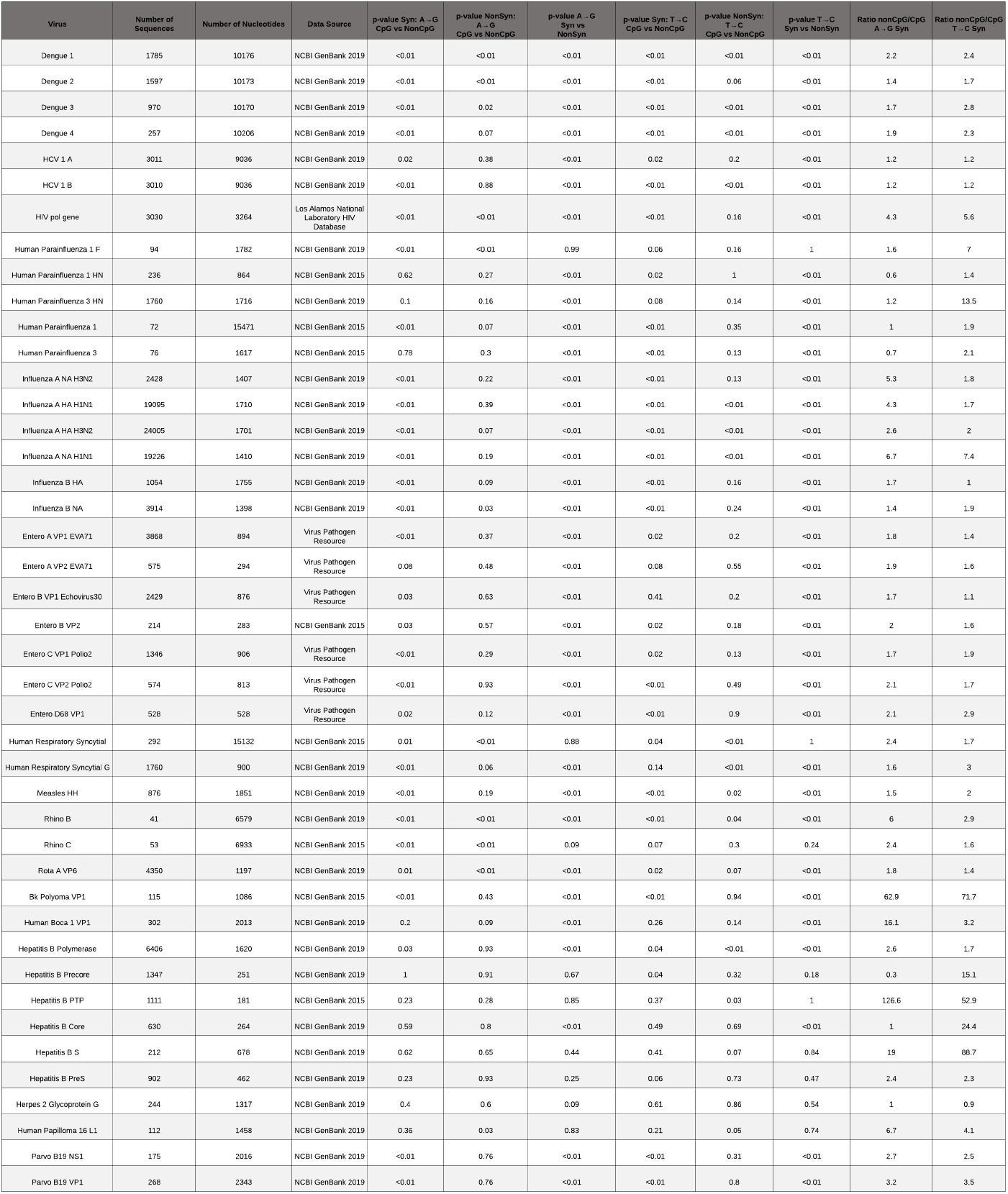
Summary of Data. All information pertaining to the datasets, such as virus name, how much and where data was collected, statistical and mathematical results.

Our study focused on transition mutations that result in CpG sites. We focused on transition mutations because they occur at a much higher rate than tranversion mutations, and provide greater power to detect meaningful differences. There are two ways for a CpG site to be formed by a transition mutation; 1) a C precedes an A (CA) and the A mutates to a G, and 2) a T precedes a G (TG) and the T mutates to a C (see figure 1).

**Figure 1:**
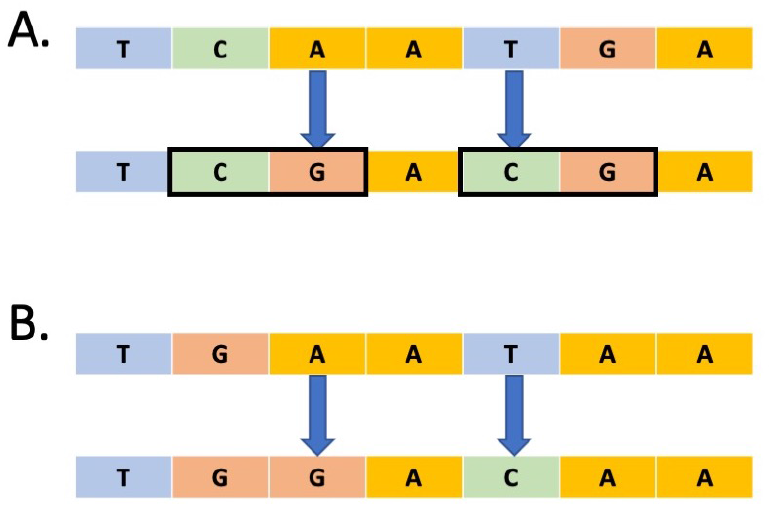
How CpG sites are created. A. There are two ways for a CpG site to be formed by a transition mutation; 1) a C precedes an A (CA) and the A mutates to a G, and 2) a T precedes a G (TG) and the T mutates to a C. B. In this study, we compare mutations that create CpG sites with similar mutations (A→G and T→C) that do not create CpG sites.

Both synonymous and non-synonymous mutations can create CpG sites. For example, when a TCA codon, which encodes Serine, mutates where the A becomes G (A→G), making the codon to TCG, this will result in a new CpG site without changing the amino acid. Comparing synonymous CpG-creating vs. synonymous non-CpG-creating mutations, we found that the frequencies of non-CpG mutations were significantly higher than those of CpG-creating mutations in 32 of the data sets (74.4%) for A→G mutations and 31 of the data sets (72.1%) for T→C mutations.

Non-synonymous mutations result in an amino acid change that alters the protein. Mutations which create a CpG site and cause a non-synonymous amino acid change are called non-synonymous CpG-creating mutations. While mutations that are non-synonymous but do not create CpG sites are called non-synonymous non-CpG-creating mutations. When comparing non-synonymous CpG-creating vs. non-synonymous non-CpG-creating mutations, non-CpG-creating mutations had a significantly higher frequency than CpG-creating mutations 25.6% of the time for A→G mutations and 30.2 % for T→C mutations (See table 1).

From our collection of viruses, we show results from three datasets as examples (Figure 2). Only A→G and T→C mutations can form CpG sites, but here we also show C→T and G→A nucleotides as a comparison. Our results varied, they ranged from exhibiting high mutation frequencies to low mutation frequencies and significant to not significant test results. The three examples chosen show the diversity of our results.

**Figure 2:**
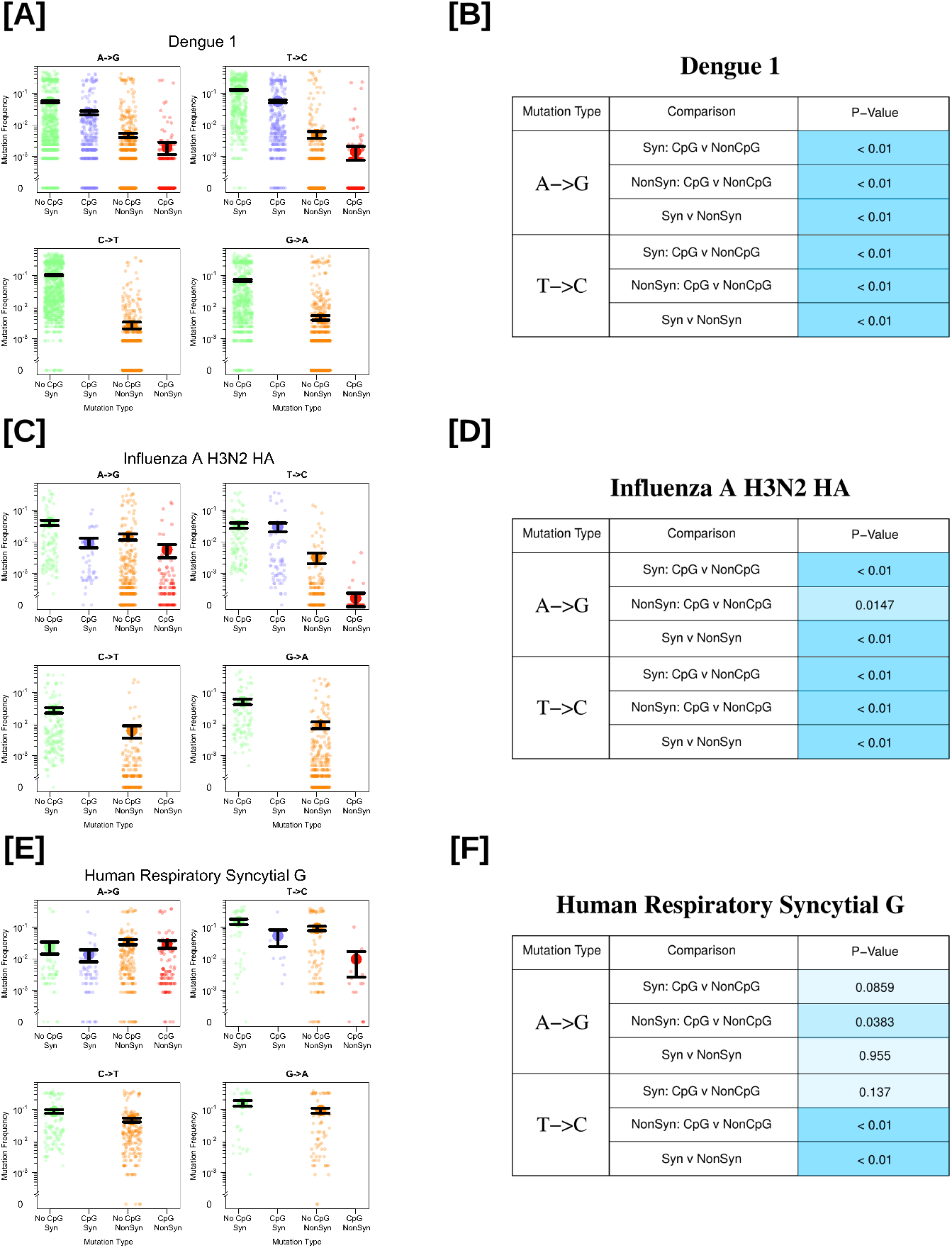
Observed transition mutation frequencies of CpG / non-CpG-creating mutations in select viral datasets (A: the whole genome of Dengue 1 virus, C: the HA gene of Influenza A virus H3N2, and E: the glycoprotein gene of Human Respiratory Syncytial virus). Each figure on the left (A, C, E) displays transition mutation frequencies, with the mean and standard errors (black lines). The Wilcoxon test results are shown on the right (B, D, F). The shade of the blue color in the P-value cell represents the significance level; darker the shade, the more significant the results are (<0.01 dark blue, 0.01-0.05 medium blue, >0.05 light blue).

In each graph, four categories are compared with one another: synonymous non-CpG-creating mutations (green), synonymous CpG-creating mutations (blue), non-synonymous non-CpG-creating mutations (orange), and non-synonymous CpG-creating mutations (red). Each colored point is the mutation frequency observed at a single position within each of these categories, along with the mean value and standard error bars (one standard error above and below the mean) in black.

Fig 2 A shows mutation frequencies for Dengue 1. Dengue’s genome is comprised of one large polyprotein. For Dengue 1, we have 1,785 sequences and 10,176 nucleotides, making this a particularly good dataset. We show frequencies of all 10,176 sites in the genome, split into the four different transition mutations (A→G, T→C, C→T, G→A) and split into synonymous(green and blue) and non-synonymous (orange and red). Non-CpG-creating mutations are green and orange, while CpG-creating mutations are red and blue. For this data set, all tested comparisons are significantly different. Synonymous CpG-creating mutations occur at lower frequencies than synonymous non-CpG-creating mutations, for both A→G and T→C mutations (green vs blue and orange vs red respectively). There is also a significant difference between the synonymous and non-synonymous mutations for both A→G and T→C mutations.

Next, we show mutation frequencies for the HA gene of the Influenza A H3N2 strain (Fig. 2 C and D). The p-values show that non-CpG-creating mutations occur at higher frequencies than CpG-creating mutations for synonymous and non-synonymous A→G and T→C mutations. For the synonymous T→C mutations, the graph shows that the mean frequencies are almost the same, but the non-parametric Wilcoxon test still detects a significant difference (p<0.01) (Fig. 2 D). The difference in frequencies between synonymous and non-synonymous mutations is significant for both A→G and T→C mutations.

Next, we show the results for Human Respiratory Syncytial Virus G gene (Fig. 2 E and F). Here there is a surprising result that for A→G mutations, we do not detect a difference between synonymous and non-synonymous mutations, whereas for T→C mutations we do. This could indicate that there is a lot of positive or balancing selection going on or that we do not have a lot of power to detect differences for this virus. The mutation frequencies of both the synonymous A→G and T→C CpG and non-CpG groups were not statistically different. For the non-synonymous mutations, on the other hand, comparisons for both A→G and T→C were significant and non-CpG-creating mutations occurred at higher frequencies (Fig. 2 F).

### Cost of CpG-creating mutations across all datasets

With a Wilcoxon test, we could determine whether CpG-creating mutations occur at lower frequencies than otherwise similar non-CpG-creating mutations, but it does not give us a sense of the effect size of this effect. To get a better sense of how much less frequent CpG-creating mutations are (and thus roughly how much more costly) we divided the mean frequency of non-CpG-creating mutations by the mean frequency of CpG-creating mutations for each of the datasets (Fig. 3). We graphed only the synonymous mutations as they more often showed a significant CpG effect.

**Figure 3:**
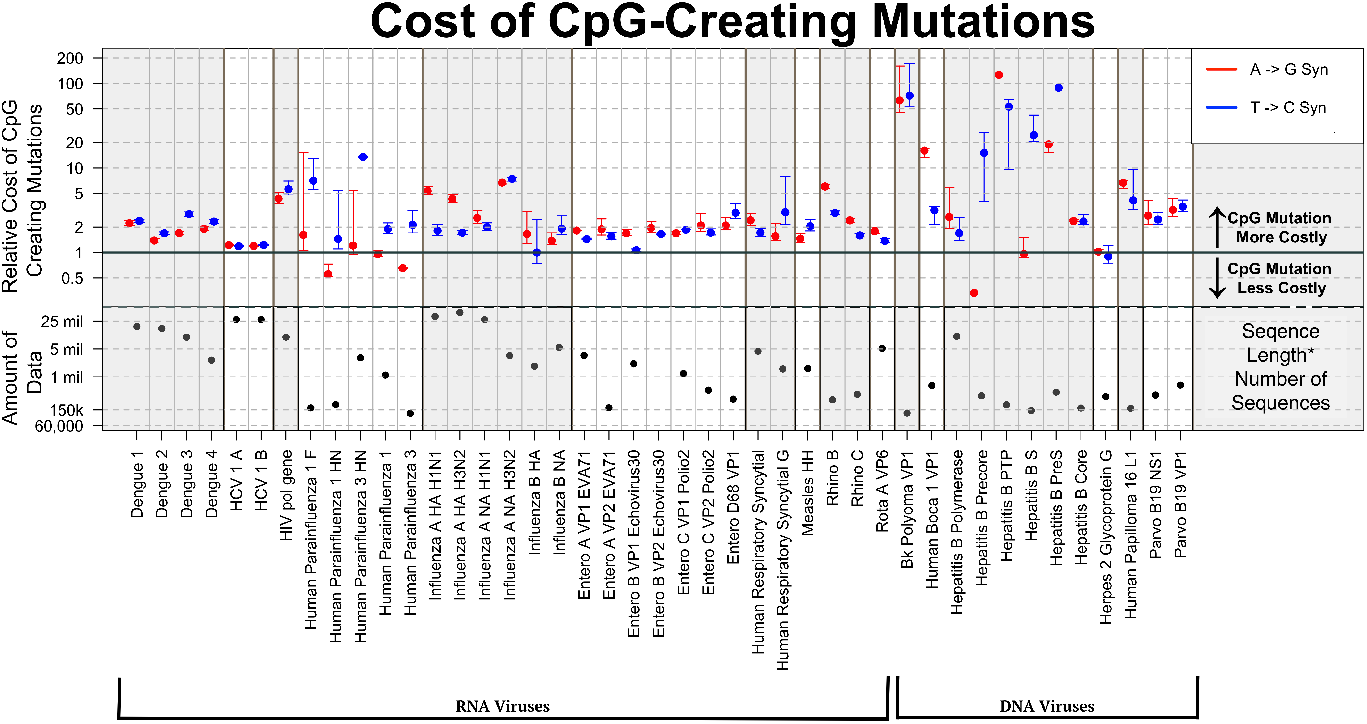
Overview of the cost associated with CpG-creating mutations. Each dot represents a ratio of the average virus mutation frequency of non-CpG-creating mutations to the average frequency of CpG-creating mutations. The bottom half of the figure depicts the total amount of data in each virus data set(the number of sequences × the number of nucleotides)

We calculated two ratios for each dataset: 1) the ratio of the mean frequency of synonymous, A→G, non-CpG-creating mutations and synonymous, A→G, CpG-creating mutations (red), and 2) the ratio of the mean frequency of synonymous, T→C, non-CpG-creating mutations and synonymous, T→C, CpG-creating mutations (blue). When these ratios are above 1 it means that the non-CpG-creating mutations have a higher average frequency than CpG-creating mutations, which shows that the CpG-creating mutations are more costly. The higher the frequency, the higher the cost of CpG-creating mutations relative to the cost of non-CpG-creating mutations. The black line in the figure 3 indicates the *ratio* = 1. Most, though not all, viruses analyzed show ratios higher than 1 (above the solid black line).

In figure 3 the viruses are arranged by genus, with RNA viruses on the left and DNA viruses on the right. We see that the calculated frequency ratios are consistently above 1 for Dengue 1-4, Hepatitis C, HIV, Influenza A and B, the Entero viruses including Polio, Human Respiratory Syncytial virus, Measles, Rhino viruses, Rota A virus, BK polyoma, Human Boca, Human papiloma and Parvo virus. Results are mixed for Parainfluenza, Hepatitis B and Herpes virus.

There is a pattern among groups of viruses where one type of mutation is more costly than the other. In Dengue and Human Parainfluenza CpG-creating T→C mutations are relatively more costly than CpG-creating A→G mutations. In Entero and Hepatitis B, on the other hand CpG-creating A→G mutations are more costly than CpG-creating T→C mutations. It is unclear whether this is an artifact of our dataset or a real effect.

Since we suspect that the amount of data available per dataset may affect our results, we plotted the product of the number of sequences and the number of nucleotides per dataset at the bottom of figure 3. In a separate figure, we also show how the amount of available data affects whether we find significant results. Figure 4A shows the comparison synonymous CpG-creating vs synonymous non-CpG-creating mutations. Each dot represents a dataset. The dots are colored by whether the two Wilcoxon tests (for A→G and T→C mutations) were both significant (blue), one was significant and one not (green) or neither was significant (red). The figure shows that, in general, having more data makes it more likely to find one or two significant results. Figure 4B shows the comparison non-synonymous CpG-creating vs non-synonymous non-CpG-creating mutations. In this case, it seems that one needs at least 2000 nucleotides or 2000 sequences to find a significant result. Finally, figure 4C shows the comparison synonymous vs non-synonymous mutations.

**Figure 4:**
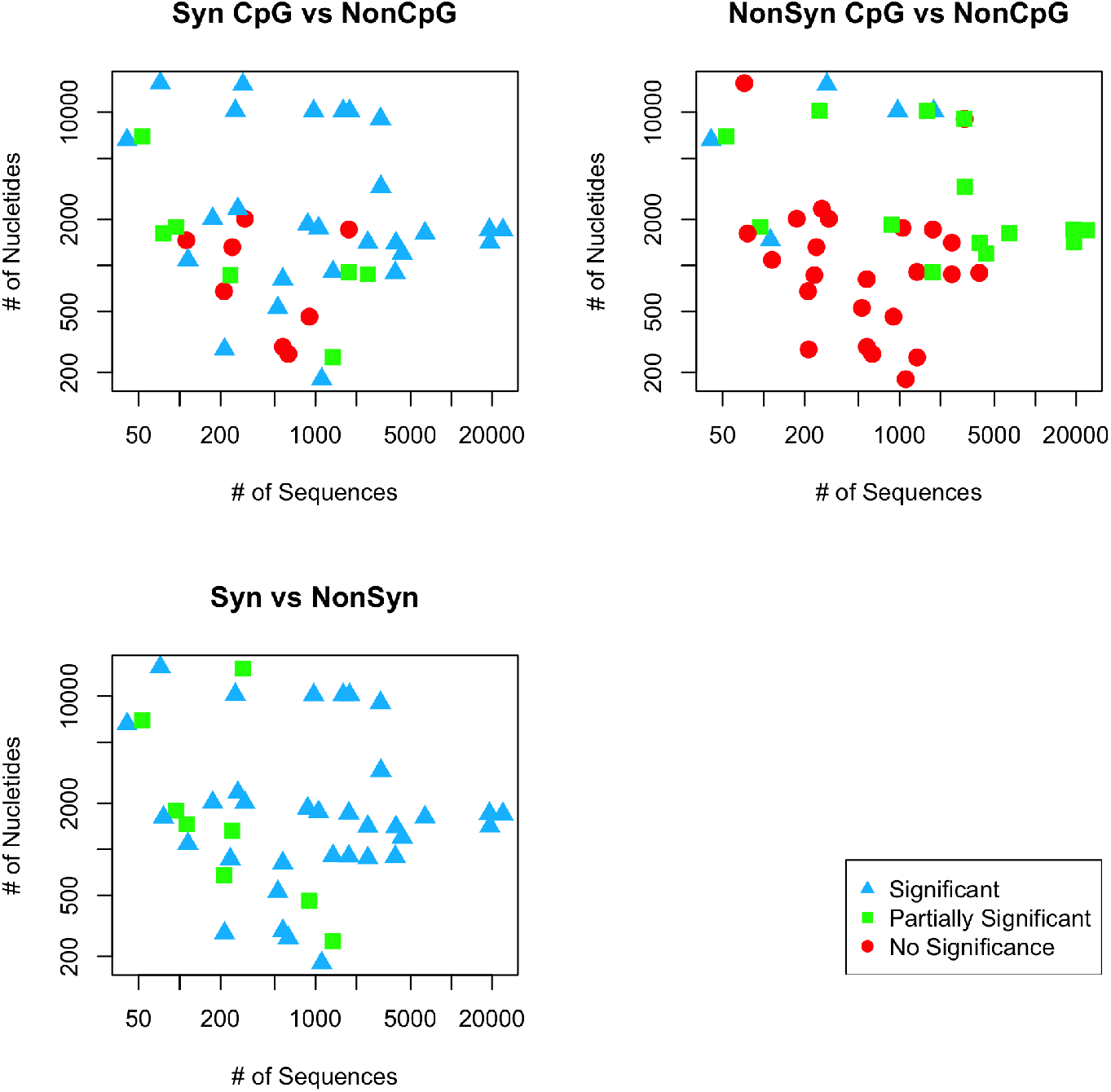
Each point represents the amount of sequences vs number of nucleotides that each data set contained. The colors and shapes represent what was found significant in each Wilcoxon test; blue triangles if both A→G and T→C are significant, green squares if only one was significant and red circles if both are not significant.

## Discussion

### CpG-creating mutations are costly in most viruses

There is previous evidence that CpG-creating mutations are costly for viruses such as HIV and Polio (Theys et al., 2018; Stern et al., 2017). It is expected that such mutations are costly in other viruses too, because CpG sites are rare in many other viruses too (Karlin and Cardon, 1994). Here we used global data for 43 viral datasets to test whether CpG sites are costly for most human viruses. For many viruses, information on within-host diversity is not readily available, so we focused, instead, on between-host diversity, using datasets with one viral sequence per patient. We expect that mutation frequencies in such datasets are determined by mutation rates, selection coefficients and stochastic effects such as drift and selective sweeps (Hartl and Clark, 2007). Our main assumption here is that stochastic effects and mutation rates affect CpG-creating and non-CpG-creating mutations equally. This means that any significant difference in mutation frequencies between CpG-creating and non-CpG-creating mutations will be due to differences in selection coefficients, which allows us to determine whether CpG-creating mutations are generally more costly than non-CpG-creating mutations (Theys et al., 2018).

We found that indeed, in the majority of viruses we tested, the mutation frequencies were significantly different between CpG-creating and non-CpG-creating mutations, which shows that there is a fitness cost to CpG-creating mutations in most viruses. We found a significant effect of CpG-creating mutations in 74.5 % of datasets for synonymous A-G mutations and in 72% of synonymous T-C mutations.

To test the statistical power of our novel approach, we also tested whether we could detect a difference in frequencies between synonymous and non-synonymous mutations. We used the same datasets and methods to demonstrate that synonymous mutations occur at higher frequencies than non-synonymous mutations. We detected a significant difference between non-synonymous mutations and synonymous mutations in 81.4 % of datasets for A→G mutations and 81.4% of datasets for T→C mutations. Hence, we detect the CpG effect almost as often as the effect of non-synonymous mutations, this despite the fact that the synonymous/non-synonymous comparison has more data to work with. The cost of CpG mutations should probably be considered near ubiquitous in human viruses.

We also tested for an effect of CpG-creating mutations among non-synonymous mutations, but found that this effect was only detected in 25.6% of datasets for A→G mutations and 30.2% of datasets for T→C mutations. One reason for this low number of significant results is probably that many non-synonymous mutations occur at very low frequencies (see figure 1A, 1C, 1E).

### Quantifying the cost

After we found that a majority of viruses displayed a lower frequency of CpG-creating mutations when compared to non-CpG-creating mutations we moved on to quantify this cost. We did this separately for A→G and T→C mutations. For each of these two types of mutations, we calculated the ratio between the mean frequency of synonymous CpG-creating mutations and the mean frequency of synonymous non-CpG-creating mutations. We hypothesize that when CpG-mutations come with a large cost, they will be found at much lower frequencies, whereas if they come with a small cost, their frequencies will only be slightly lower than those of non-CpG-creating mutations. Therefore, the ratio we calculate will give us a sense of the relative cost of CpG sites in different viruses.

The levels of the cost ratio vary widely between viruses, with some clear differences between viral genera. For example, for HIV, the ratio is near 5 for both AG and CT mutations. This shows that CpG sites in HIV come with a large cost, as shown before based on a different data set(Theys et al., 2018). On the other hand, in Hepatitis C the ratio is close to 1 for both genotype 1A and 1B. The Wilcoxon tests were significant for Hepatitis C, but the fact that the ratio is close to 1 shows that the effect size is small. We find similar results when we look at within-host diversity for HCV using another dataset (Tisthammer, unpublished). The cost ratio for Bk Polyoma is very high, but the amount of data for this virus is quite low and we expect that with more data, the ratio will go down to a level similar to the other viruses.

We find more variable cost ratios in the DNA viruses than in the RNA viruses. This may be because of the smaller sample sizes for DNA viruses, or it may be that different selection pressures are at play in DNA viruses versus RNA viruses. In RNA viruses, we expect that the mammalian immune system recognizes CpG sites and forces the viruses to mimic the low CpG content in mammalian genomes (Takata et al., 2017). In DNA viruses, it is not clear if the same mechanism is at work, though unmethylated CpG sites are expected to stimulate the immune response (Hoelzer et al., 2008).

The cost ratio was calculated for both A→G and C→T mutations. These two ratios are not necessarily equal. In some viruses, we see surprising patterns in the cost ratios. For example, in the Dengue viruses, T→C CpG-creating mutations (blue) are relatively more costly than A→G mutations (red). In Influenza A however, the trend is in the other direction, where T→C CpG-creating mutations (blue) are relatively less costly than A→G mutations (red). Further studies are needed to determine what causes these patterns.

### Limitations and future studies

We only included datasets with at least 60,000 data points per dataset (number of nucleotides * number of sequences). However, we still find that our larger datasets are more likely to yield significant results (Figure 4). This suggests that increasing either the number of sequences or the sequence length for some of the viral datasets will increase the number of datasets with significant results.

Another limitation of our study is that we used one sequence per patient. For some viruses, such as HIV and HCV, it is possible to use within-patient genetic diversity to study costs of mutations within the host (Wang et al., 2010; Rambaut et al., 2004; Alizon et al., 2011). This is possible for these viruses because patients are infected for a long time and there is an expectation that mutation and selection occur within the host. For other viruses, it is not clear whether it is possible to study within-host fitness costs separately from between-host effects. If patients are infected with a diverse sample of the virus, then within-host mutation and selection may not be the dominant effects that shape within-host genetic diversity (Varble et al., 2014; Poon et al., 2016). For those types of viruses, studying within-host and between-host diversity may lead to the same results, and having data on within-host diversity may not increase our knowledge of fitness costs of mutations.

In conclusion, we find that CpG-creating mutations are costly for most human viruses. For viruses in which we do not detect an effect of CpG-creating mutations, it is likely because of a small sample size. Future work should focus on better understanding why the cost of CpG-creating mutations is higher in some viruses than others.

## Methods

### Data and R scripts

Data and R scripts are available on Github: https://github.com/Vcaudill/SomethingCool-CpGSites-.git

### Data Collection

The sequences were retrieved from the NCBI Genbank, the HIV Databases (http://www.hiv.lanl.gov/), and the Virus Pathogen resource (VPR, https://www.viprbrc.org/) using R scripts or manually. We selected viral sequences from a human host, and for proteins required for viral fitness (e.g. VP1, VP2, envelope protein). Dengue, Entero, and Polio sequences were all collected through the VPR, HIV sequences from the HIV Database, and HCV, Human Parainfluenza, Influenza, Human Respiratory Syncytial, Measles, Rhino, Rota, BK, Human Boca, Hepatitis B, Human Heperies, Human Papilloma, and Parvo from Genbank.

### Further Data Preparation and Filtering

After data collection, obtained sequences were aligned and trimmed using Geneious v.11.1.4. After checking the alignment, an online translation tool (SIB Web Team, 1993) was used to identify coding regions. Once a coding region was found, the sequences were verified using NCBI BLAST. Upon verification, consensus sequences for each virus/protein data set were generated using R or Geneious. A custom R script was also used to identify stop codons created by mutations in the coding sequences. All stop codons and insertions were removed from the alignment. Accurate estimation of mutation frequencies requires sufficient data points. Therefore, we calculated data points as the number of sequences multiplied by the number of nucleotides, and removed data sets that had less than 60,000 data points. We were able to collect sufficient data for 43 data sets.

### Data Analysis

For each of the 43 data sets, the consensus sequence was translated to create a wild type protein sequence. For each nucleotide, we determined whether a transition mutation would change the amino acid and/or create a CpG site. We determined whether the transition mutation was synonymous, non-synonymous or nonsense by comparing the wild type amino acid to the mutated amino acid. We calculated the frequency of the transition mutation for each nucleotide in the data set by dividing the number of observed transition mutations by the sum of the number of transition mutations and the wild type nucleotide.

### Statistical Analysis

To determine if CpG sites were costly to viruses, the data were separated into groups. First, the sites were split into four categories; each represented a consensus nucleotide and its transition mutated form (Adenine to Guanine (A→G), Thymine to Cytosine (T→C), Cytosine to Thymine (C→T), or Guanine to Adenine (G→A)). The nucleotides were then sectioned into groups of synonymous and non-synonymous, and further by CpG-creating or non-CpG-creating mutations. (Fig. 5). A Wilcoxon rank-sum test was performed to determine if the mutation frequencies differed between groups of synonymous *vs*. non-synonymous, and CpG *vs*. non-CpG-creating mutations (Fig. 2). To calculate a “cost ratio” of CpG-creating transition mutations, we divided the mean mutation frequency of non-CpG-creating mutations by CpG-creating mutations of the same type (Fig. 3).

**Figure 5:**
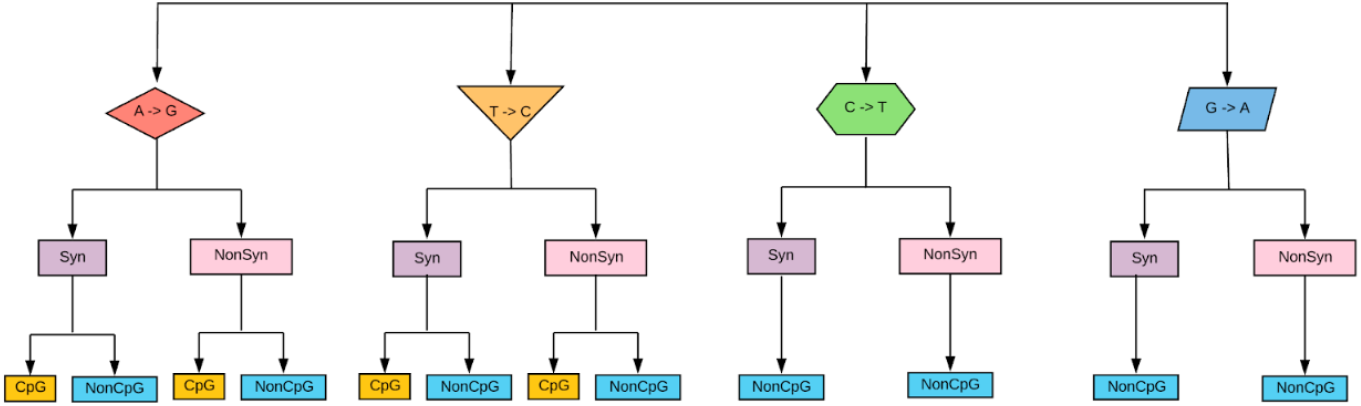
A pictorial representation of 12 mutation groups. Each wildtype nucleotide was categorized into synonymous or non-synonymous by the resulting amino acid. For A and T, we further separated the groups into CpG or non-CpG-creating mutations (Nucleotides C and G cannot form CpG sites).

## Supporting information

Supplemental Results

## Acknowledgements

We thank Adi Stern and the Stern lab for discussion. Pleuni Pennings, Victoria Caudill, Sarina Qin, Ryan Winstead, Jasmeen Kaur, Esteban Geo Pineda, Emily Fryer, Kaho Tisthammer and Anjani Pradhananga were supported by NSF grant # 1655212 to Pleuni S. Pennings. Sarah Cobey and Pleuni Pennings were supported by a grant from the National Evolutionary Synthesis Center (NESCent), NSF # EF-0905606. Angeline K. Chemel, Edward Hopson and Nicole S. Rodriques were supported by the NIH MARC grant (T34-GM008574). Dwayne Evans was supported by the NIH MA/MS-PhD Bridge grant (R25-GM048972). Dwayne Evans, Alejandro G Lopez, A.R. Mahoney, Rebecca L. Melton and Ryan Winstead were supported by the NIH RISE grant (R25-GM059298) Angeline K. Chemel and Kellen Hopp were supported by the NSF STC CCC grant (DBI 1548297). Dwayne Evans and Alejandro G Lopez were supported by a Genentech Foundation MS Dissertation Scholarship.

## References

Alizon, S., Luciani, F., and Regoes, R. R. (2011). Epidemiological and clinical consequences of within-host evolution. Trends in microbiology, 19(1):24–32

Burns, C. C., Campagnoli, R., Shaw, J., Vincent, A., Jorba, J., and Kew, O. (2009). Genetic inactivation of poliovirus infectivity by increasing the frequencies of CpG and UpA dinucleotides within and across synonymous capsid region codons. J. Virol., 83(19):9957–9969.

Cuevas, J. M., Geller, R., Garijo, R., Lopez-Aldeguer, J., and Sanjuan, R. (2015). Extremely High Mutation Rate of HIV-1 In Vivo. PLoS Biol., 13(9):e1002251.

Duffy, S. (2018). Why are rna virus mutation rates so damn high? PLOS Biology, 16(8):1–6.

Hartl, D. L. and Clark, A. G. (2007). Principles of population genetics.

Hoelzer, K., Shackelton, L. A., and Parrish, C. R. (2008). Presence and role of cytosine methylation in dna viruses of animals. Nucleic acids research, 36(9):2825–2837.

Karlin, S. and Cardon, L. R. (1994). Computational dna sequence analysis. Annual Review of Microbiology, 48(1):619–654. PMID: 7826021.

Murphy, K. M. and Weaver, C. (2016). Janeway’s Immunobiology. Garland Science, Taylor Francis Group, LLC.

Notkins, A. L., Mergenhagen, S. E., and Howard, R. J. (1970). Effect of virus infections on the function of the immune system. Annual Review of Microbiology, 24(1):525–538. PMID: 4928348.

Poon, L. L., Song, T., Rosenfeld, R., Lin, X., Rogers, M. B., Zhou, B., Sebra, R., Halpin, R. A., Guan, Y., Twaddle, A., et al. (2016). Quantifying influenza virus diversity and transmission in humans. Nature genetics, 48(2):195.

Rambaut, A., Posada, D., Crandall, K. A., and Holmes, E. C. (2004). The causes and consequences of hiv evolution. Nature Reviews Genetics, 5(1):52.

Sanjuán, R., Moya, A., and Elena, S. F. (2004). The distribution of fitness effects caused by single-nucleotide substitutions in an rna virus. Proceedings of the National Academy of Sciences, 101(22):8396–8401.

SIB Web Team (1993). Sib expasy bioformatics resources portal. https://web.expasy.org/translate/.

Stern, A., Doron-Faigenboim, A., Erez, E., Martz, E., Bacharach, E., and Pupko, T. (2007). Selecton 2007: advanced models for detecting positive and purifying selection using a bayesian inference approach. Nucleic acids research, 35(suppl 2):W506–W511.

Stern, A., Te Yeh, M., Zinger, T., Smith, M., Wright, C., Ling, G., Nielsen, R., Macadam, A., and Andino, R. (2017). The evolutionary pathway to virulence of an rna virus. Cell, 169(1):35–46.

Takata, M. A., Goncalves-Carneiro, D., Zang, T. M., Soll, S. J., York, A., Blanco-Melo, D., and Bieniasz, P. D. (2017). CG dinucleotide suppression enables antiviral defence targeting non-self RNA. Nature, 550(7674):124–127.

Theys, K., Feder, A. F., Gelbart, M., Hartl, M., Stern, A., and Pennings, P. S. (2018). Correction: Within-patient mutation frequencies reveal fitness costs of CpG dinucleotides and drastic amino acid changes in HIV. PLoS Genet., 14(12):e1007855.

Varble, A., Albrecht, R. A., Backes, S., Crumiller, M., Bouvier, N. M., Sachs, D., García-Sastre, A., et al. (2014). Influenza a virus transmission bottlenecks are defined by infection route and recipient host. Cell host & microbe, 16(5):691–700.

Wang, G. P., Sherrill-Mix, S. A., Chang, K.-M., Quince, C., and Bushman, F. D. (2010). Hepatitis c virus transmission bottlenecks analyzed by deep sequencing. Journal of virology, 84(12):6218–6228.

Zanini, F., Puller, V., Brodin, J., Albert, J., and Neher, R. A. (2017). In vivo mutation rates and the landscape of fitness costs of hiv-1. Virus evolution, 3(1):vex003.

