## Supplemental Results for "CpG-creating Mutations are Costly in Many Human Viruses"

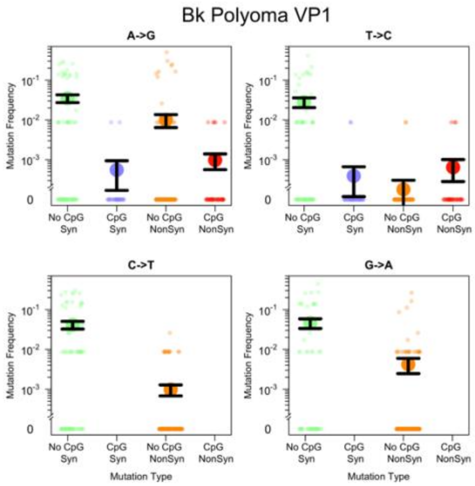


### Bk Polyoma VP1

| Mutation Type | Comparison | P−Value |
|---|---|---|
| A−>G | Syn: CpG v NonCpG | < 0.01 |
| | NonSyn: CpG v NonCpG | 0.43 |
| | Syn v NonSyn | < 0.01 |
| T−>C | Syn: CpG v NonCpG | < 0.01 |
| | NonSyn: CpG v NonCpG | 0.937 |
| | Syn v NonSyn | < 0.01 |

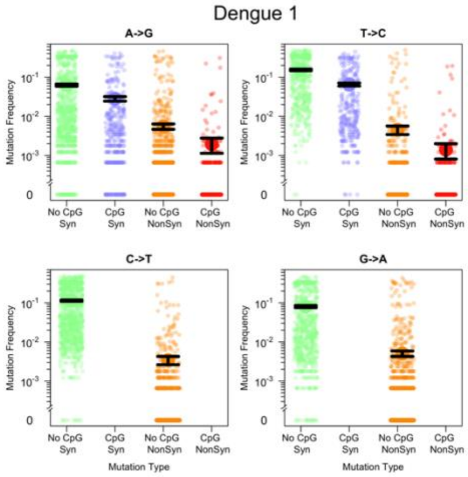


### Dengue 1

| Mutation Type | Comparison | P−Value |
|---|---|---|
| A−>G | Syn: CpG v NonCpG | < 0.01 |
| | NonSyn: CpG v NonCpG | < 0.01 |
| | Syn v NonSyn | < 0.01 |
| T−>C | Syn: CpG v NonCpG | < 0.01 |
| | NonSyn: CpG v NonCpG | < 0.01 |
| | Syn v NonSyn | < 0.01 |

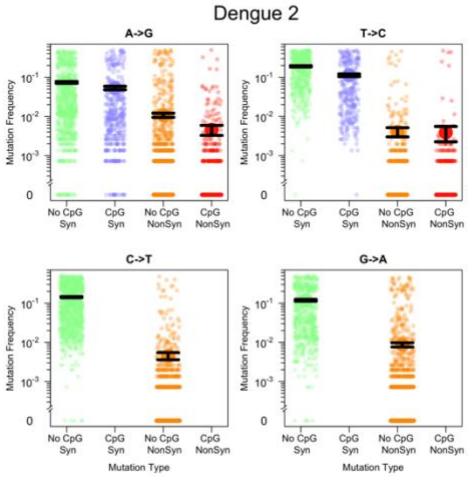


### Dengue 2

| Mutation Type | Comparison | P−Value |
|---|---|---|
| A−>G | Syn: CpG v NonCpG | < 0.01 |
| | NonSyn: CpG v NonCpG | < 0.01 |
| | Syn v NonSyn | < 0.01 |
| T−>C | Syn: CpG v NonCpG | < 0.01 |
| | NonSyn: CpG v NonCpG | 0.061 |
| | Syn v NonSyn | < 0.01 |

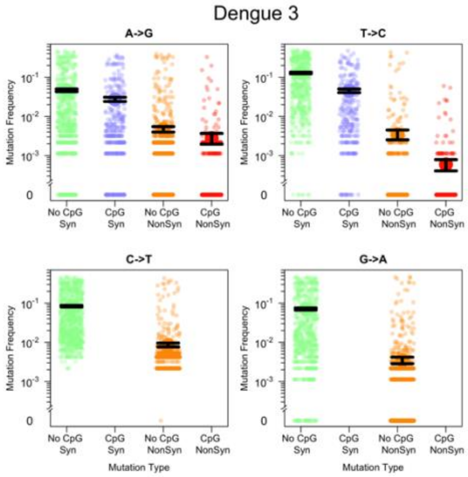


### Dengue 3

| Mutation Type | Comparison | P−Value |
|---|---|---|
| A−>G | Syn: CpG v NonCpG | < 0.01 |
| | NonSyn: CpG v NonCpG | 0.0227 |
| | Syn v NonSyn | < 0.01 |
| T−>C | Syn: CpG v NonCpG | < 0.01 |
| | NonSyn: CpG v NonCpG | < 0.01 |
| | Syn v NonSyn | < 0.01 |

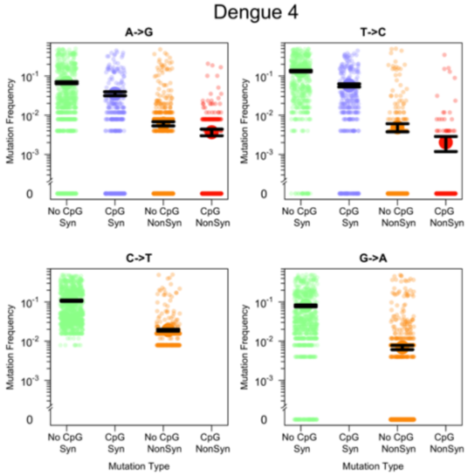


### Dengue 4

| Mutation Type | Comparison | P−Value |
|---|---|---|
| A−>G | Syn: CpG v NonCpG | < 0.01 |
| | NonSyn: CpG v NonCpG | 0.073 |
| | Syn v NonSyn | < 0.01 |
| T−>C | Syn: CpG v NonCpG | < 0.01 |
| | NonSyn: CpG v NonCpG | < 0.01 |
| | Syn v NonSyn | < 0.01 |

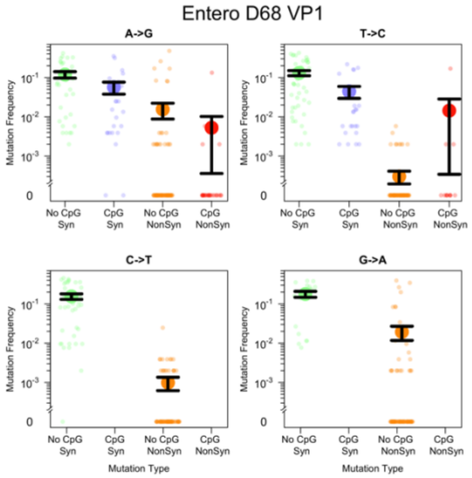


### Entero D68 VP1

| Mutation Type | Comparison | P−Value |
|---|---|---|
| A−>G | Syn: CpG v NonCpG | 0.0158 |
| | NonSyn: CpG v NonCpG | 0.12 |
| | Syn v NonSyn | < 0.01 |
| T−>C | Syn: CpG v NonCpG | < 0.01 |
| | NonSyn: CpG v NonCpG | 0.897 |
| | Syn v NonSyn | < 0.01 |

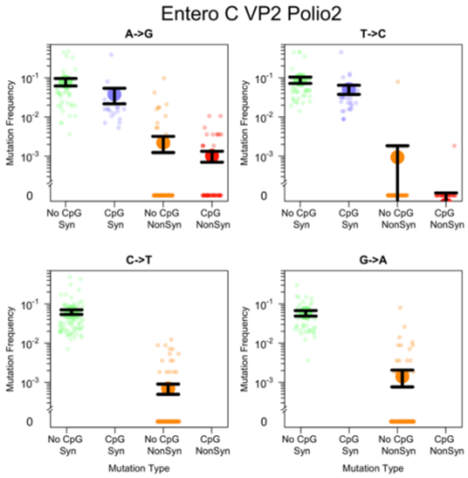


### Entero C VP2 Polio2

| Mutation Type | Comparison | P−Value |
|---|---|---|
| A−>G | Syn: CpG v NonCpG | < 0.01 |
| | NonSyn: CpG v NonCpG | 0.926 |
| | Syn v NonSyn | < 0.01 |
| T−>C | Syn: CpG v NonCpG | < 0.01 |
| | NonSyn: CpG v NonCpG | 0.488 |
| | Syn v NonSyn | < 0.01 |

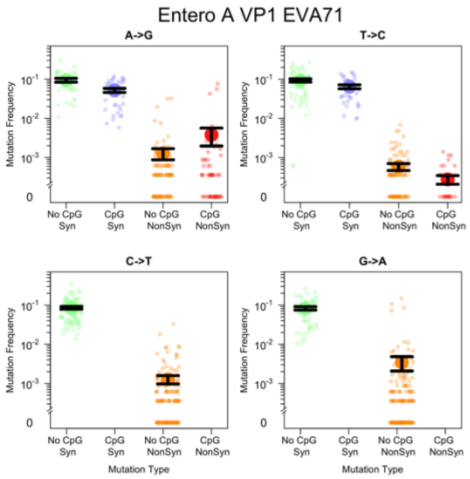


### Entero A VP1 EVA71

| Mutation Type | Comparison | P−Value |
|---|---|---|
| A−>G | Syn: CpG v NonCpG | < 0.01 |
| | NonSyn: CpG v NonCpG | 0.37 |
| | Syn v NonSyn | < 0.01 |
| T−>C | Syn: CpG v NonCpG | 0.0219 |
| | NonSyn: CpG v NonCpG | 0.196 |
| | Syn v NonSyn | < 0.01 |

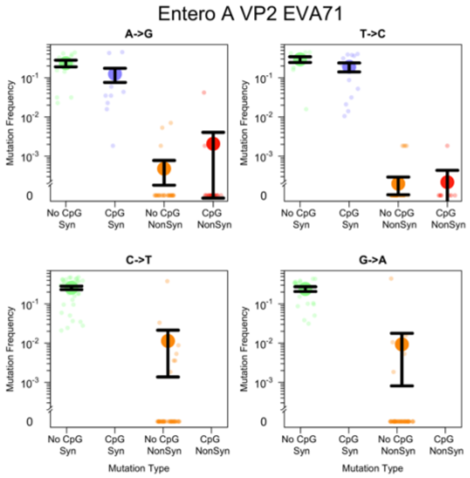


#### Entero A VP2 EVA71

| Mutation Type | Comparison | P−Value |
|---|---|---|
| A−>G | Syn: CpG v NonCpG | 0.0783 |
| | NonSyn: CpG v NonCpG | 0.477 |
| | Syn v NonSyn | < 0.01 |
| T−>C | Syn: CpG v NonCpG | 0.0847 |
| | NonSyn: CpG v NonCpG | 0.545 |
| | Syn v NonSyn | < 0.01 |

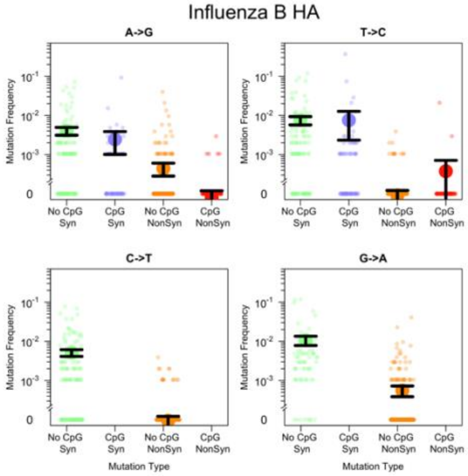


### Influenza B HA

| Mutation Type | Comparison | P-Value |
|---|---|---|
| A->G | Syn: CpG v NonCpG | < 0.01 |
| | NonSyn: CpG v NonCpG | 0.0946 |
| | Syn v NonSyn | < 0.01 |
| T->C | Syn: CpG v NonCpG | < 0.01 |
| | NonSyn: CpG v NonCpG | 0.162 |
| | Syn v NonSyn | < 0.01 |

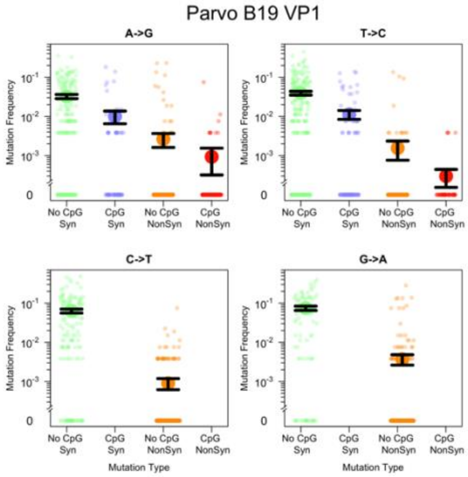


#### Parvo B19 VP1

| Mutation Type | Comparison | P−Value |
|---|---|---|
| A−>G | Syn: CpG v NonCpG | < 0.01 |
| | NonSyn: CpG v NonCpG | 0.755 |
| | Syn v NonSyn | < 0.01 |
| T−>C | Syn: CpG v NonCpG | < 0.01 |
| | NonSyn: CpG v NonCpG | 0.801 |
| | Syn v NonSyn | < 0.01 |

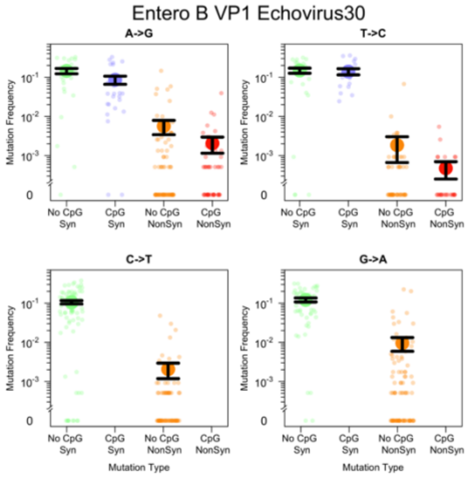


#### Entero B VP1 Echovirus30

| Mutation Type | Comparison | P–Value |
|---|---|---|
| A–>G | Syn: CpG v NonCpG | 0.0256 |
| | NonSyn: CpG v NonCpG | 0.625 |
| | Syn v NonSyn | < 0.01 |
| T–>C | Syn: CpG v NonCpG | 0.407 |
| | NonSyn: CpG v NonCpG | 0.204 |
| | Syn v NonSyn | < 0.01 |

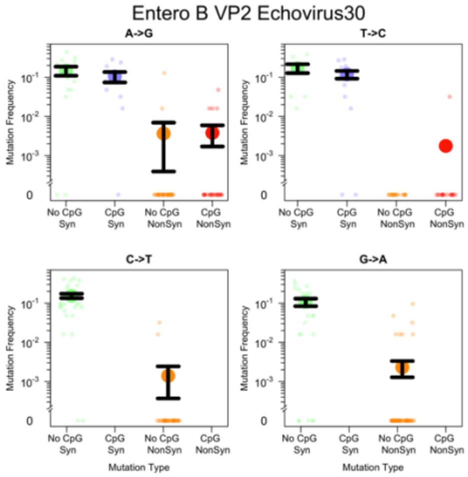


### Entero B VP2 Echovirus30

| Mutation Type | Comparison | P–Value |
|---|---|---|
| A–>G | Syn: CpG v NonCpG | 0.256 |
| | NonSyn: CpG v NonCpG | 0.922 |
| | Syn v NonSyn | < 0.01 |
| T–>C | Syn: CpG v NonCpG | 0.135 |
| | NonSyn: CpG v NonCpG | 0.866 |
| | Syn v NonSyn | < 0.01 |

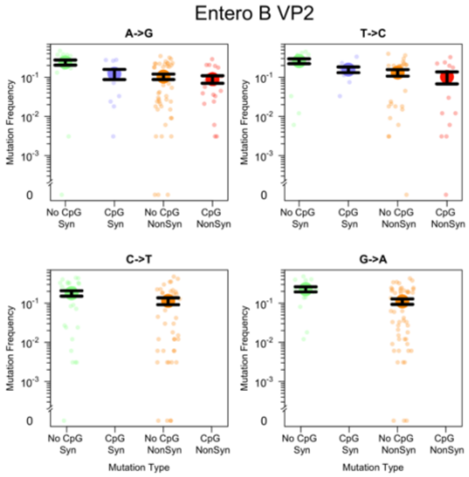


#### Entero B VP2 Echovirus30

| Mutation Type | Comparison | P−Value |
| --- | --- | --- |
| A−>G | Syn: CpG v NonCpG | 0.0336 |
| | NonSyn: CpG v NonCpG | 0.567 |
| | Syn v NonSyn | < 0.01 |
| T−>C | Syn: CpG v NonCpG | 0.02 |
| | NonSyn: CpG v NonCpG | 0.177 |
| | Syn v NonSyn | < 0.01 |

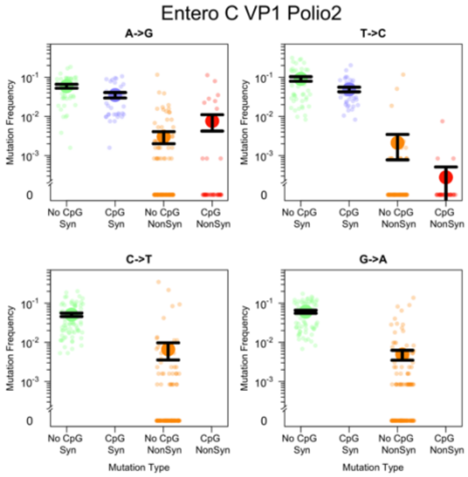


#### Entero C VP1 Polio2

| Mutation Type | Comparison | P−Value |
|---|---|---|
| A−>G | Syn: CpG v NonCpG | < 0.01 |
| | NonSyn: CpG v NonCpG | 0.288 |
| | Syn v NonSyn | < 0.01 |
| T−>C | Syn: CpG v NonCpG | 0.0244 |
| | NonSyn: CpG v NonCpG | 0.126 |
| | Syn v NonSyn | < 0.01 |

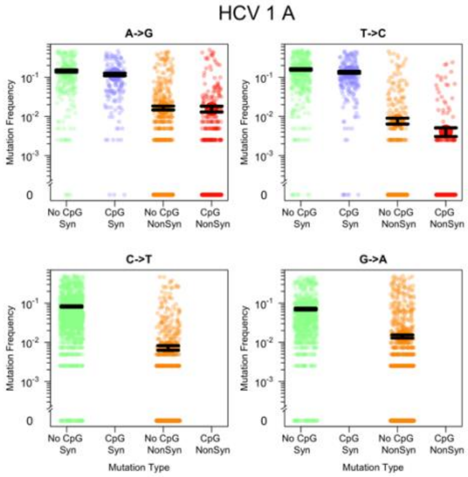


#### HCV 1 A

| Mutation Type | Comparison | P−Value |
|---|---|---|
| A−>G | Syn: CpG v NonCpG | 0.0209 |
| | NonSyn: CpG v NonCpG | 0.364 |
| | Syn v NonSyn | < 0.01 |
| T−>C | Syn: CpG v NonCpG | 0.0271 |
| | NonSyn: CpG v NonCpG | 0.178 |
| | Syn v NonSyn | < 0.01 |

HCV 1 B

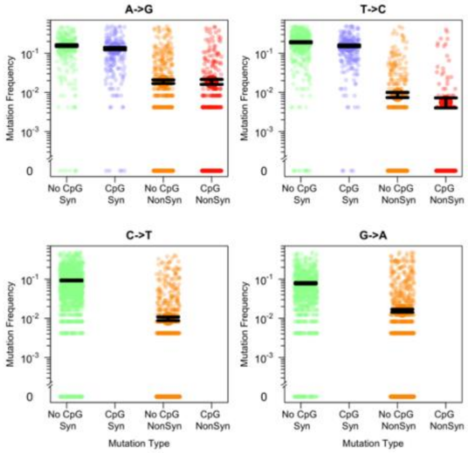


#### HCV 1 B

| Mutation Type | Comparison | P−Value |
|---|---|---|
| A−>G | Syn: CpG v NonCpG | < 0.01 |
| | NonSyn: CpG v NonCpG | 0.883 |
| | Syn v NonSyn | < 0.01 |
| T−>C | Syn: CpG v NonCpG | < 0.01 |
| | NonSyn: CpG v NonCpG | < 0.01 |
| | Syn v NonSyn | < 0.01 |

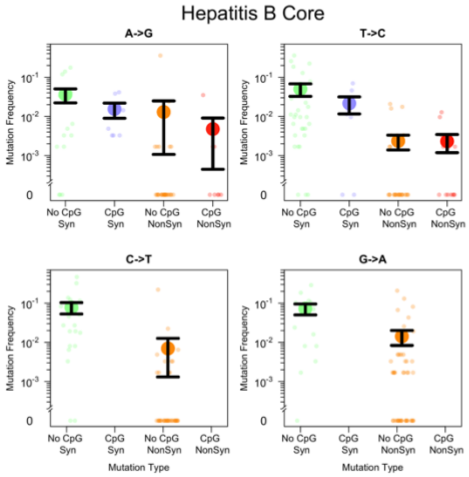


#### Hepatitis B Core

| Mutation Type | Comparison | P–Value |
|---|---|---|
| A–>G | Syn: CpG v NonCpG | 0.593 |
| | NonSyn: CpG v NonCpG | 0.798 |
| | Syn v NonSyn | < 0.01 |
| T–>C | Syn: CpG v NonCpG | 0.492 |
| | NonSyn: CpG v NonCpG | 0.692 |
| | Syn v NonSyn | < 0.01 |

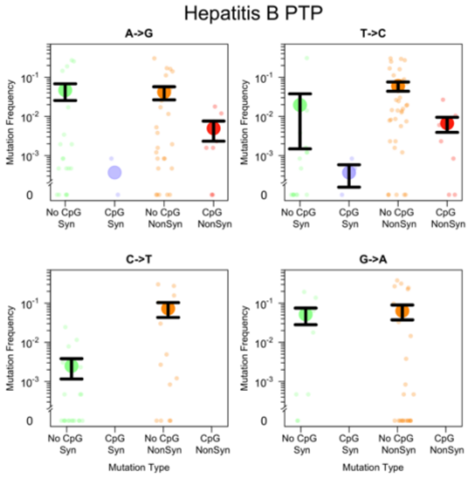


#### Hepatitis B PTP

| Mutation Type | Comparison | P–Value |
|---|---|---|
| A–>G | Syn: CpG v NonCpG | 0.232 |
| | NonSyn: CpG v NonCpG | 0.276 |
| | Syn v NonSyn | 0.85 |
| T–>C | Syn: CpG v NonCpG | 0.369 |
| | NonSyn: CpG v NonCpG | 0.0277 |
| | Syn v NonSyn | 1 |

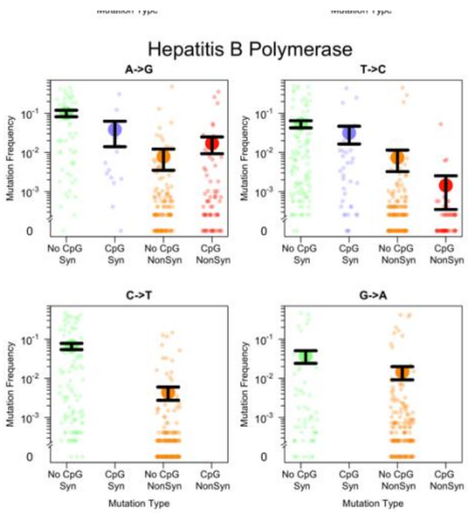


### Hepatitis B Polymerase

| Mutation Type | Comparison | P−Value |
|---|---|---|
| A−>G | Syn: CpG v NonCpG | 0.0335 |
| | NonSyn: CpG v NonCpG | 0.93 |
| | Syn v NonSyn | < 0.01 |
| T−>C | Syn: CpG v NonCpG | 0.0433 |
| | NonSyn: CpG v NonCpG | < 0.01 |
| | Syn v NonSyn | < 0.01 |

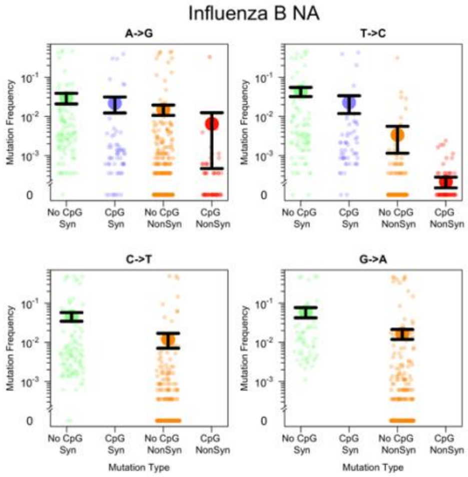


**Influenza B NA**

| Mutation Type | Comparison | P−Value |
|---|---|---|
| A−>G | Syn: CpG v NonCpG | < 0.01 |
| | NonSyn: CpG v NonCpG | 0.0328 |
| | Syn v NonSyn | < 0.01 |
| T−>C | Syn: CpG v NonCpG | < 0.01 |
| | NonSyn: CpG v NonCpG | 0.237 |
| | Syn v NonSyn | < 0.01 |

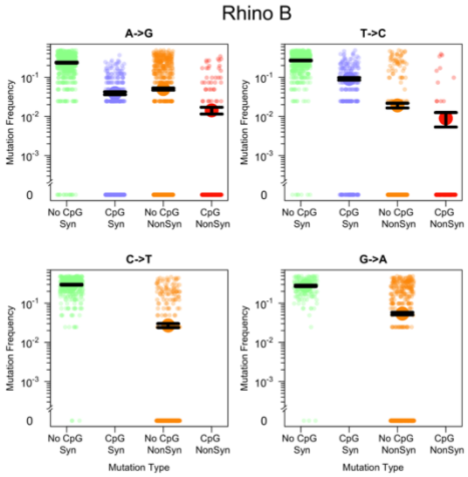


### Rhino B

| Mutation Type | Comparison | P–Value |
|---|---|---|
| A–>G | Syn: CpG v NonCpG | < 0.01 |
| | NonSyn: CpG v NonCpG | < 0.01 |
| | Syn v NonSyn | < 0.01 |
| T–>C | Syn: CpG v NonCpG | < 0.01 |
| | NonSyn: CpG v NonCpG | 0.0426 |
| | Syn v NonSyn | < 0.01 |

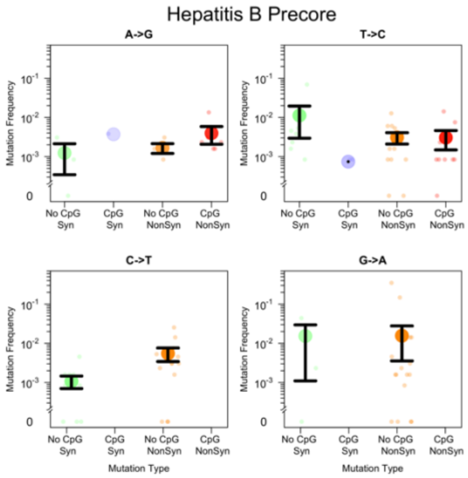


### Hepatitis B Precore

| Mutation Type | Comparison | P−Value |
|---|---|---|
| A−>G | Syn: CpG v NonCpG | 1 |
| | NonSyn: CpG v NonCpG | 0.908 |
| | Syn v NonSyn | 0.667 |
| T−>C | Syn: CpG v NonCpG | 0.0429 |
| | NonSyn: CpG v NonCpG | 0.316 |
| | Syn v NonSyn | 0.175 |

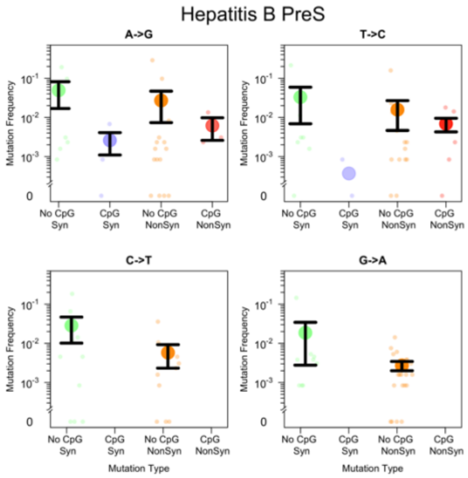


### Hepatitis B PreS

| Mutation Type | Comparison | P−Value |
|---|---|---|
| A−>G | Syn: CpG v NonCpG | 0.226 |
| | NonSyn: CpG v NonCpG | 0.933 |
| | Syn v NonSyn | 0.249 |
| T−>C | Syn: CpG v NonCpG | 0.0575 |
| | NonSyn: CpG v NonCpG | 0.727 |
| | Syn v NonSyn | 0.466 |

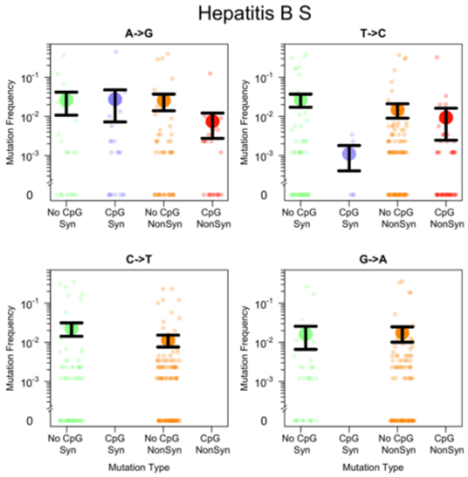


#### Hepatitis B S

| Mutation Type | Comparison | P−Value |
|---|---|---|
| A−>G | Syn: CpG v NonCpG | 0.616 |
| | NonSyn: CpG v NonCpG | 0.654 |
| | Syn v NonSyn | 0.441 |
| T−>C | Syn: CpG v NonCpG | 0.412 |
| | NonSyn: CpG v NonCpG | 0.0733 |
| | Syn v NonSyn | 0.838 |

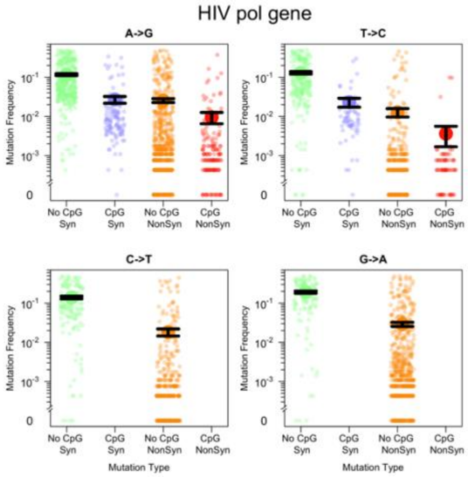


### HIV pol gene

| Mutation Type | Comparison | P−Value |
|---|---|---|
| A−>G | Syn: CpG v NonCpG | < 0.01 |
| | NonSyn: CpG v NonCpG | < 0.01 |
| | Syn v NonSyn | < 0.01 |
| T−>C | Syn: CpG v NonCpG | < 0.01 |
| | NonSyn: CpG v NonCpG | 0.155 |
| | Syn v NonSyn | < 0.01 |

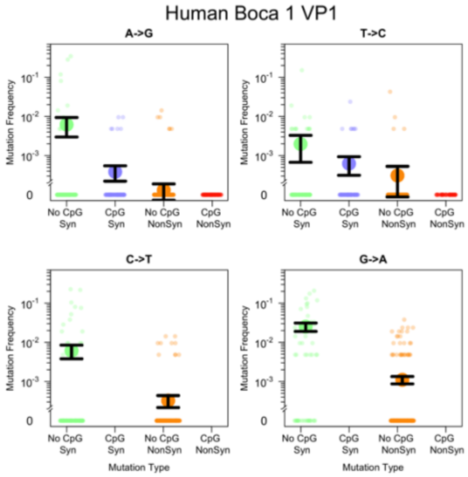


### Human Boca 1 VP1

| Mutation Type | Comparison | P–Value |
|---|---|---|
| A–>G | Syn: CpG v NonCpG | 0.195 |
| | NonSyn: CpG v NonCpG | 0.0873 |
| | Syn v NonSyn | < 0.01 |
| T–>C | Syn: CpG v NonCpG | 0.258 |
| | NonSyn: CpG v NonCpG | 0.141 |
| | Syn v NonSyn | < 0.01 |

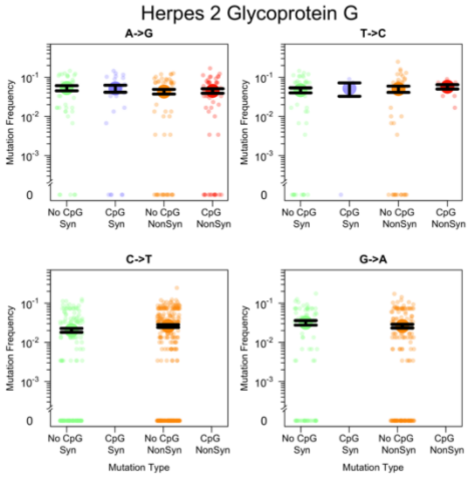


### Herpes 2 Glycoprotein G

| Mutation Type | Comparison | P–Value |
|---|---|---|
| A–>G | Syn: CpG v NonCpG | 0.398 |
| | NonSyn: CpG v NonCpG | 0.602 |
| | Syn v NonSyn | 0.0905 |
| T–>C | Syn: CpG v NonCpG | 0.609 |
| | NonSyn: CpG v NonCpG | 0.857 |
| | Syn v NonSyn | 0.544 |

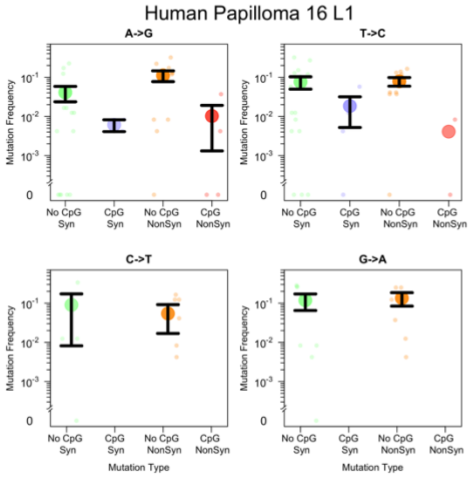


### Human Papilloma 16 L1

| Mutation Type | Comparison | P−Value |
|---|---|---|
| A−>G | Syn: CpG v NonCpG | 0.361 |
| | NonSyn: CpG v NonCpG | 0.0321 |
| | Syn v NonSyn | 0.826 |
| T−>C | Syn: CpG v NonCpG | 0.21 |
| | NonSyn: CpG v NonCpG | 0.0483 |
| | Syn v NonSyn | 0.742 |

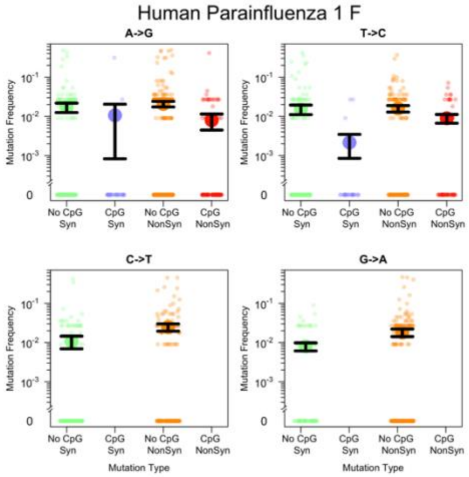


### Human Parainfluenza 1 F

| Mutation Type | Comparison | P−Value |
|---|---|---|
| A−>G | Syn: CpG v NonCpG | < 0.01 |
| | NonSyn: CpG v NonCpG | < 0.01 |
| | Syn v NonSyn | 0.988 |
| T−>C | Syn: CpG v NonCpG | 0.0559 |
| | NonSyn: CpG v NonCpG | 0.159 |
| | Syn v NonSyn | 1 |

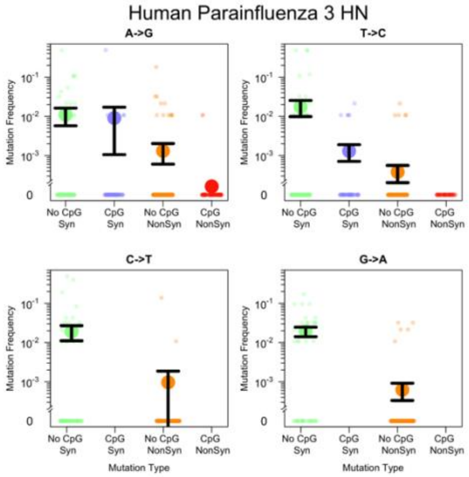


### Human Parainfluenza 3 HN

| Mutation Type | Comparison | P−Value |
|---|---|---|
| A−>G | Syn: CpG v NonCpG | 0.103 |
| | NonSyn: CpG v NonCpG | 0.157 |
| | Syn v NonSyn | < 0.01 |
| T−>C | Syn: CpG v NonCpG | 0.0772 |
| | NonSyn: CpG v NonCpG | 0.139 |
| | Syn v NonSyn | < 0.01 |

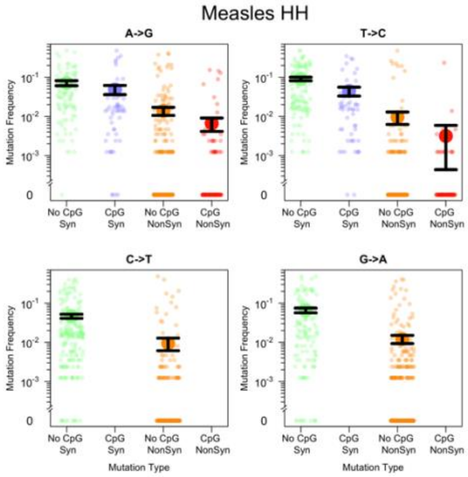


### Measles HH

| Mutation Type | Comparison | P−Value |
|---|---|---|
| A−>G | Syn: CpG v NonCpG | < 0.01 |
| | NonSyn: CpG v NonCpG | 0.193 |
| | Syn v NonSyn | < 0.01 |
| T−>C | Syn: CpG v NonCpG | < 0.01 |
| | NonSyn: CpG v NonCpG | 0.0174 |
| | Syn v NonSyn | < 0.01 |

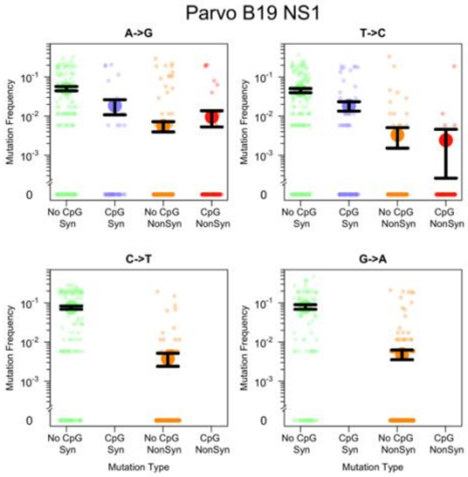


#### Parvo B19 NS1

| Mutation Type | Comparison | P−Value |
|---|---|---|
| A−>G | Syn: CpG v NonCpG | < 0.01 |
| | NonSyn: CpG v NonCpG | 0.763 |
| | Syn v NonSyn | < 0.01 |
| T−>C | Syn: CpG v NonCpG | < 0.01 |
| | NonSyn: CpG v NonCpG | 0.308 |
| | Syn v NonSyn | < 0.01 |

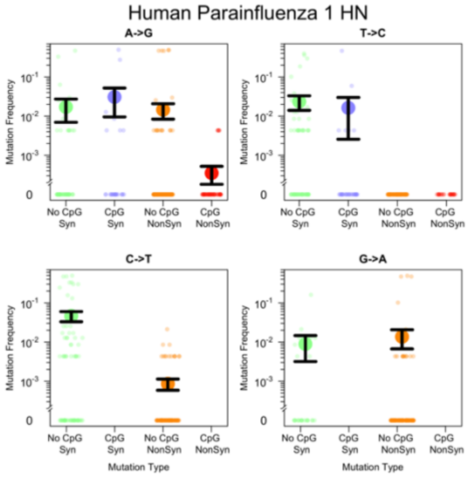


### Human Parainfluenza 1 HN

| Mutation Type | Comparison | P–Value |
|---|---|---|
| A–>G | Syn: CpG v NonCpG | 0.608 |
| | NonSyn: CpG v NonCpG | 0.3 |
| | Syn v NonSyn | < 0.01 |
| T–>C | Syn: CpG v NonCpG | 0.0176 |
| | NonSyn: CpG v NonCpG | 1 |
| | Syn v NonSyn | < 0.01 |

Human Parainfluenza 1

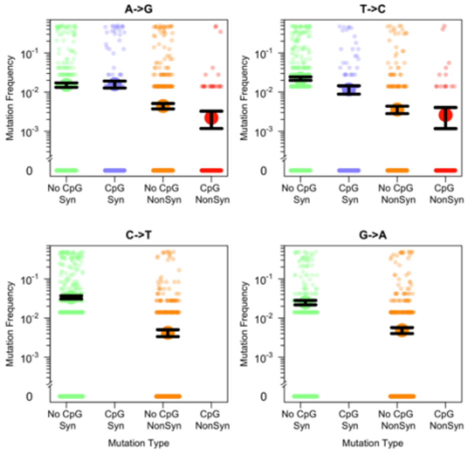


### Human Parainfluenza 1

| Mutation Type | Comparison | P−Value |
|---|---|---|
| A−>G | Syn: CpG v NonCpG | < 0.01 |
| | NonSyn: CpG v NonCpG | 0.0664 |
| | Syn v NonSyn | < 0.01 |
| T−>C | Syn: CpG v NonCpG | < 0.01 |
| | NonSyn: CpG v NonCpG | 0.35 |
| | Syn v NonSyn | < 0.01 |

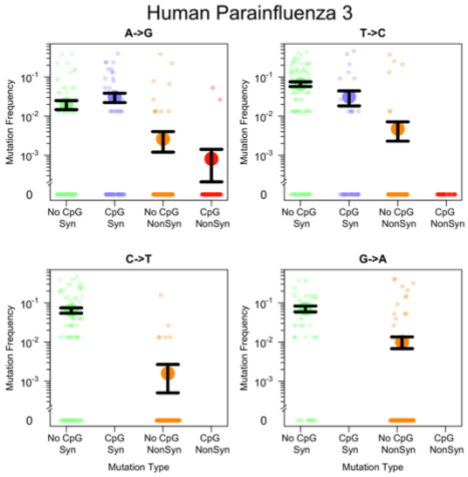


### Human Parainfluenza 3

| Mutation Type | Comparison | P−Value |
|---|---|---|
| A−>G | Syn: CpG v NonCpG | 0.777 |
| | NonSyn: CpG v NonCpG | 0.297 |
| | Syn v NonSyn | < 0.01 |
| T−>C | Syn: CpG v NonCpG | < 0.01 |
| | NonSyn: CpG v NonCpG | 0.132 |
| | Syn v NonSyn | < 0.01 |

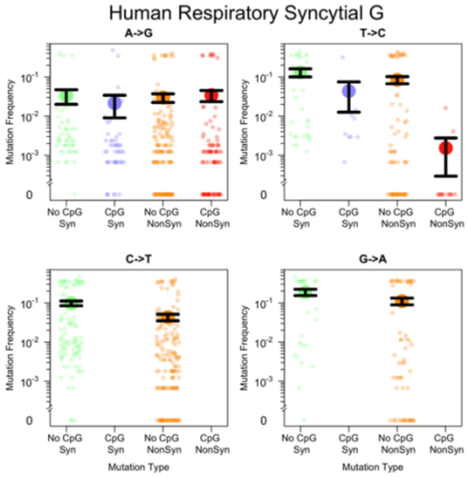


### Human Respiratory Syncytial G

| Mutation Type | Comparison | P−Value |
|---|---|---|
| A−>G | Syn: CpG v NonCpG | < 0.01 |
| | NonSyn: CpG v NonCpG | 0.056 |
| | Syn v NonSyn | < 0.01 |
| T−>C | Syn: CpG v NonCpG | 0.143 |
| | NonSyn: CpG v NonCpG | < 0.01 |
| | Syn v NonSyn | < 0.01 |

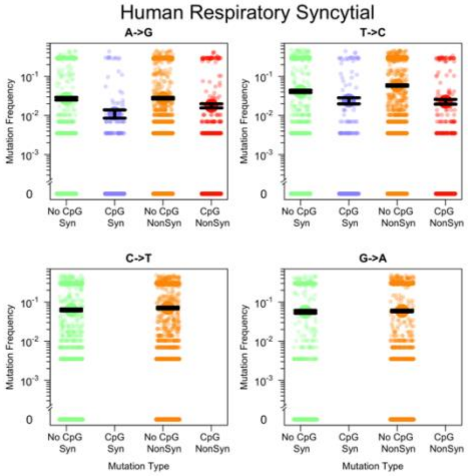


### Human Respiratory Syncytial

| Mutation Type | Comparison | P−Value |
|---|---|---|
| A−>G | Syn: CpG v NonCpG | 0.0115 |
| | NonSyn: CpG v NonCpG | < 0.01 |
| | Syn v NonSyn | 0.881 |
| T−>C | Syn: CpG v NonCpG | 0.0363 |
| | NonSyn: CpG v NonCpG | < 0.01 |
| | Syn v NonSyn | 1 |

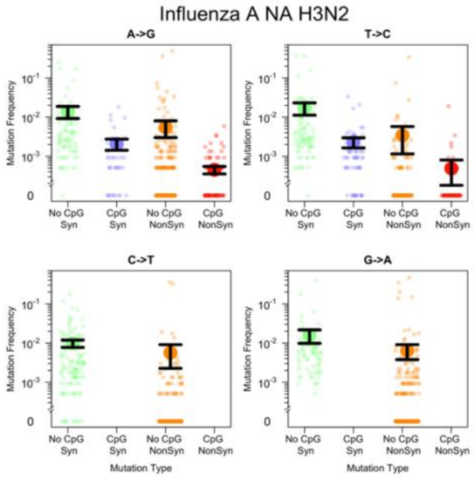


### Influenza A NA H3N2

| Mutation Type | Comparison | P−Value |
|---|---|---|
| A−>G | Syn: CpG v NonCpG | < 0.01 |
| | NonSyn: CpG v NonCpG | 0.224 |
| | Syn v NonSyn | < 0.01 |
| T−>C | Syn: CpG v NonCpG | < 0.01 |
| | NonSyn: CpG v NonCpG | 0.126 |
| | Syn v NonSyn | < 0.01 |

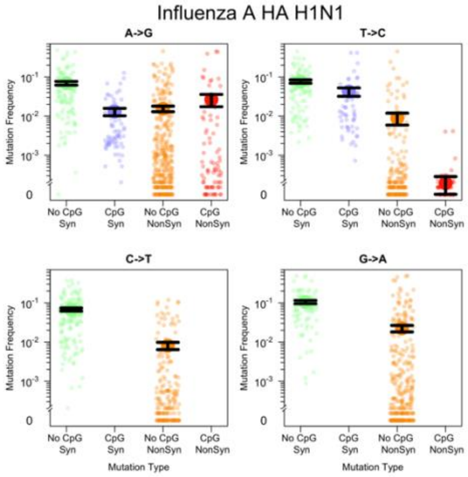


### Influenza A HA H1N1

| Mutation Type | Comparison | P−Value |
|---|---|---|
| A−>G | Syn: CpG v NonCpG | < 0.01 |
| | NonSyn: CpG v NonCpG | 0.385 |
| | Syn v NonSyn | < 0.01 |
| T−>C | Syn: CpG v NonCpG | < 0.01 |
| | NonSyn: CpG v NonCpG | < 0.01 |
| | Syn v NonSyn | < 0.01 |

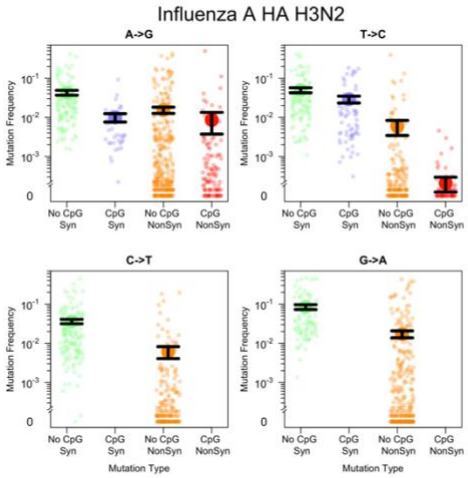


### Influenza A HA H3N2

| Mutation Type | Comparison | P−Value |
|---|---|---|
| A−>G | Syn: CpG v NonCpG | < 0.01 |
| | NonSyn: CpG v NonCpG | 0.0677 |
| | Syn v NonSyn | < 0.01 |
| T−>C | Syn: CpG v NonCpG | < 0.01 |
| | NonSyn: CpG v NonCpG | < 0.01 |
| | Syn v NonSyn | < 0.01 |

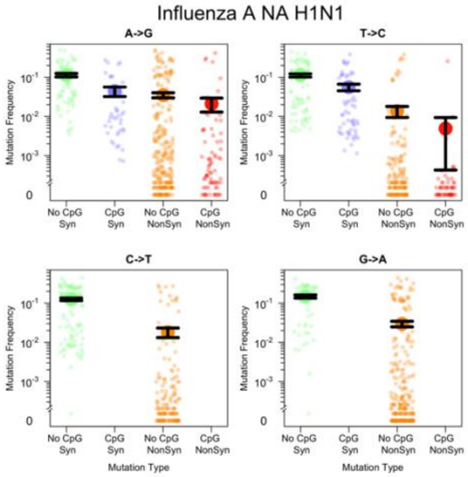


### Influenza A NA H1N1

| Mutation Type | Comparison | P−Value |
|---|---|---|
| A−>G | Syn: CpG v NonCpG | < 0.01 |
| | NonSyn: CpG v NonCpG | 0.19 |
| | Syn v NonSyn | < 0.01 |
| T−>C | Syn: CpG v NonCpG | < 0.01 |
| | NonSyn: CpG v NonCpG | < 0.01 |
| | Syn v NonSyn | < 0.01 |

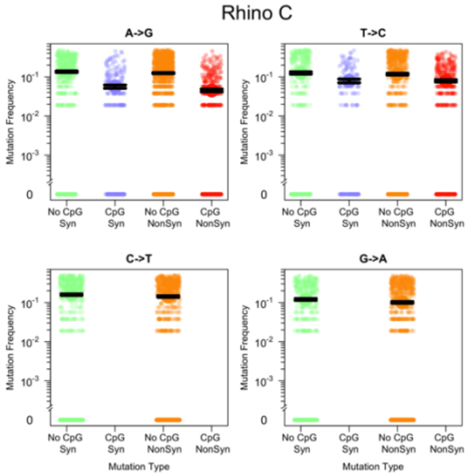


### Rhino C

| Mutation Type | Comparison | P−Value |
|---|---|---|
| A−>G | Syn: CpG v NonCpG | < 0.01 |
| | NonSyn: CpG v NonCpG | < 0.01 |
| | Syn v NonSyn | 0.0852 |
| T−>C | Syn: CpG v NonCpG | 0.0695 |
| | NonSyn: CpG v NonCpG | 0.301 |
| | Syn v NonSyn | 0.237 |

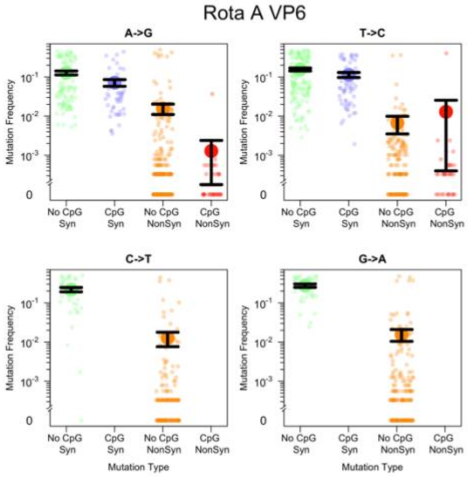


### Rota A VP6

| Mutation Type | Comparison | P−Value |
|---|---|---|
| A−>G | Syn: CpG v NonCpG | 0.0103 |
| | NonSyn: CpG v NonCpG | < 0.01 |
| | Syn v NonSyn | < 0.01 |
| T−>C | Syn: CpG v NonCpG | 0.0211 |
| | NonSyn: CpG v NonCpG | 0.0678 |
| | Syn v NonSyn | < 0.01 |
